# Duhuo Jisheng Decoction Regulates Intracellular Zinc Homeostasis by Enhancing Autophagy via PTEN/Akt/mTOR to Improve Knee Cartilage Degeneration

**DOI:** 10.1101/2023.08.19.553976

**Authors:** Ye-Hui Wang, Yi Zhou, Xiang Gao, Sheng Sun, Yi-Zhou Xie, You-Peng Hu, Yang Fu, Xiao-Hong Fan, Quan Xie

**Author notes:** Corresponding authors at: No.37, Shi’er Qiao Road, Jinniu District, Chengdu 610075, Sichuan Province, China. E-mail addresses (Xiao-Hong Fan), (Quan Xie). Ye-Hui Wang and Yi Zhou contributed equally to the work.

## Abstract

**Background:** Articular cartilage degeneration as well as cartilage matrix degradation is one of the key pathological changes in the early stage of knee osteoarthritis (KOA). However, currently, there are limited early prevention and treatment options available. Duhuo Jisheng Decoction (DHJSD) is a formula from *Bei Ji Qian jin Yao Fang* compiled by Sun Simiao in the Tang Dynasty of China. As a complementary therapy, it is widely used to treat early-stage KOA in China, but its mechanism has not been fully elucidated.

**Objective:** This study is aiming at investigating the potential role and mechanism of DHJSD in protecting cartilage from degradation.

**Methods:** The mechanism of DHJSD in alleviating OA was explored by gene silencing technology combined with a series of functional experiments in primary rat chondrocytes. Next, 25 wistar rats were used to validate the results obtained *in vitro*. The PTEN, Akt, mTOR, MMP13, Zn, collagen II, autophagy and apoptosis were determined.

**Results:** DHJSD reduced the phosphorylation of Akt and mTOR and the expression of zinc, MMP13, Bax and Bcl2. DHJSD increased the level of autophagy and the expression of autophagy proteins LC3 and Beclin1. After silencing PTEN gene, the phosphorylation levels of Akt and mTOR and the effects of Bax, Bcl2, LC3 and Beclin1 were weakened by DHJSD. DHJSD increased the formation of autophagosomes in chondrocytes. Histopathological staining revealed that DHJSD had a protective effect on cartilage.

**Conclusion:** DHJSD inhibits Akt/mTOR signaling pathway by targeting PTEN to promote autophagy in chondrocytes, which may be closely to repress the formation of MMP-13 by regulating the level of zinc in chondrocytes.

## 1. Introduction

Duhuo Jisheng Decoction (DHJSD) is a formula from *Beiji Qianjin Yaofang* compiled by Sun Simiao in the Tang Dynasty of China, which is composed of *Angelica pubescens* Maxim. (Duhuo, Chinese), *Asarum heterotropoides* F.Schmidt (Xixin, Chinese), *Saposhnikovia divaricata* (Turcz. ex Ledeb.) Schischk. (Fangfeng, Chinese), *Neolitsea cassia* (L.) Kosterm (Rougui, Chinese), *Gentiana macrophylla* Pall. (Qinjiao, Chinese), *Taxillus chinensis* (DC.) Danser (Sangjisheng, Chinese), *Eucommia ulmoides* Oliv. (Duzhong, Chinese), Cyathula officinalis K.C.Kuan, (Chuanniuxi, Chinese), *Paeonia lactiflora* Pall. (Baishao, Chinese), *Rehmannia glutinosa* (Gaertn.) DC. (Dihuang, Chinese), *Angelica sinensis* (Oliv.) Diels (Danggui, Chinese), *Panax ginseng* C.A.Mey. (Renshen, Chinses), *Conioselinum anthriscoides ‘Chuanxiong’* (Chuanxiong, Chinese), *Smilax glabra* Roxb. (Fuling, Chinese), *Glycyrrhiza glabra* L. (Gancao, Chinese). All the names of the plants in DHJSD have been checked with http://mpns.kew.org on June 12th, 2023. They are combined in the ratio of (9:6:6:6:6:6:6:6:6:6:6:6:6:6:6) as shown in **Table 1**. The clinical effect of DHJSD in the treatment of knee osteoarthritis is certain^[1-3]^, but there is a lack of mechanism research. Studies have shown that DHJSD can reduce the level of intracellular zinc and interfere with the degradation of cell matrix to play a protective role in chondrocytes^[4]^. The pathogenesis of OA is the result of the joint action of many factors, and the excessive degradation of extracellular matrix is one of the main pathological processes of articular cartilage degeneration^[5]^, and the degradation of chondrocytes matrix mediated by zinc is an important link in cartilage degeneration^[6, 7]^. Autophagy is closely related to intracellular zinc homeostasis^[8-10]^. Therefore, it can be speculated that autophagy is closely related to the regulation of zinc homeostasis in chondrocytes, and autophagy may be one of the upstream mechanisms of zinc-mediated cartilage degeneration.

**Table 1.**
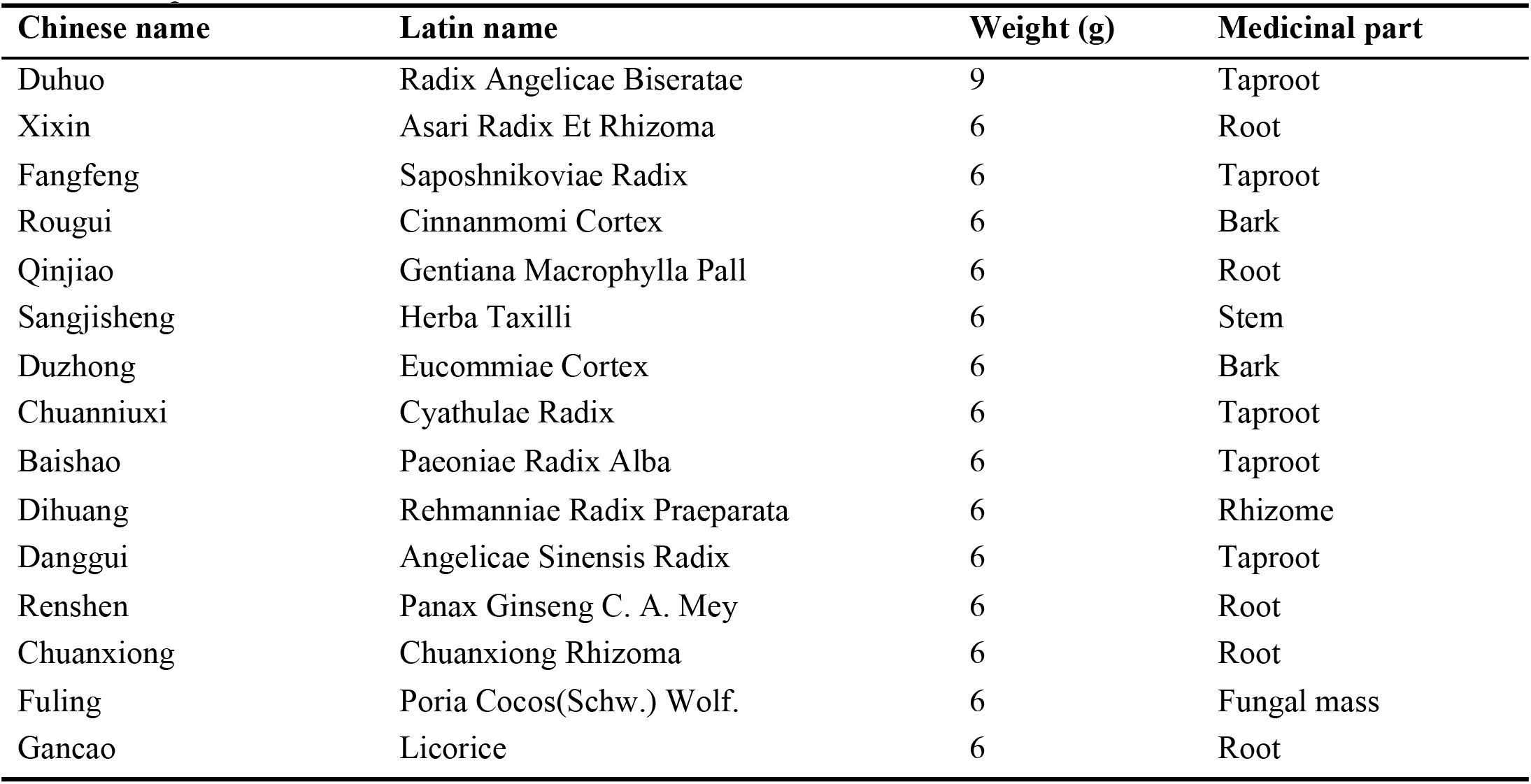
Recipe of DHJSD.

| Chinese name | Latin name | Weight (g) | Medicinal part |
| --- | --- | --- | --- |
| Duhuo | Radix Angelicae Biseratae | 9 | Taproot |
| Xixin | Asari Radix Et Rhizoma | 6 | Root |
| Fangfeng | Saposhnikoviae Radix | 6 | Taproot |
| Rougui | Cinnamomi Cortex | 6 | Bark |
| Qinjiao | Gentiana Macrophylla Pall | 6 | Root |
| Sangjiisheng | Herba Taxilli | 6 | Stem |
| Duzhong | Eucommiae Cortex | 6 | Bark |
| Chuanniuxi | Cyathulae Radix | 6 | Taproot |
| Baishao | Paeoniae Radix Alba | 6 | Taproot |
| Dihuang | Rehmanniae Radix Praeparata | 6 | Rhizome |
| Danggui | Angelicae Sinensis Radix | 6 | Taproot |
| Renshen | Panax Ginseng C. A. Mey | 6 | Root |
| Chuanxiong | Chuanxiong Rhizoma | 6 | Root |
| Fuling | Poria Cocos(Schw.) Wolf. | 6 | Fungal mass |
| Gancao | Licorice | 6 | Root |

PI3K/Akt/mTOR is an important intracellular autophagy signal regulation pathway. Activation of this pathway can regulate cell growth and metabolism, inhibit the activity of a variety of autophagy-related proteins, and reduce the ability of autophagy, which is closely related to cell growth, energy metabolism, and the occurrence and development of a variety of diseases^[11, 12]^. PI3K/Akt/mTOR signaling pathway is closely related to autophagy in chondrocytes ^[12]^. LC3, as a molecular marker of autophagy, is located in the cytoplasm. During autophagy, LC3-I binds covalently to phosphatidyleolamine to form LC3-II and localizes on the autophagosome membrane to participate in the formation of autophagosome ^[13]^. Inhibition of this pathway can increase the level of autophagy ^[14]^. Phosphatase and tensin homolog deleted on chromosome ten (PTEN) is a typical tumor suppressor gene that inhibits the PI3K/Akt/mTOR growth signaling cascade^[15]^. This pathway can be regulated through multiple sites ^[16]^, and its dysfunction will lead to the dysregulation of this pathway and other pathways, leading to overgrowth^[17]^. At present, PTEN protein has been found to have four heterogeneous structures, PTEN-L, PTEN-M, PTEN-N and PTEN-O, among which PTEN-L plays an important role in the regulation of PI3K/Akt/mTOR signaling pathway and autophagy^[18, 19]^. PTEN can directly inhibit the PI3K/Akt/mTOR pathway, thereby increasing the level of autophagy and reducing apoptosis ^[20]^. PTEN also plays an important role in the regulation of PI3K/Akt/mTOR in the metabolism of chondrocytes. Zhang et al.^[21]^ found that miRNA-132 could up-regulate PTEN and inhibit PI3K/Akt signaling pathway to enhance the level of autophagy in chondrocytes. An Gui-feng et al.^[22]^ studied human chondrocytes and found that miR-365-3p negatively regulated PTEN expression to inhibit the proliferation, migration and invasion of articular chondrocytes and promote their apoptosis. Therefore, the effect and mechanism of DHJSD on cartilage *in vitro* and *in vivo* were investigated based on PTEN/Akt/mTOR, autophagy and the zinc homeostasis in chondrocytes in this study.

## 2. Materials and methods

### 2.1 Reagents

The riboFECTTMCP(R10035.8) and siRNA(R10043.8) were supplied by RiboBio(Guangzhou, China). Animal total RNA isolation kit (RE-03014, Spec. 200) was purchased from Foregene (Chengdu, China). PrimeScript-RT reagent kit (RR047A, Spec.100) and TB Green TM Premix Ex TaqTM II (Tli RNaseH Plus) (RR820A, Spec. 200) were supplied by Takara (Beijing, China). IP cell lysate (P0013) and BCA Protein Concentration Assay Kit (P0009) were obtained from Beyotime (Shanghai, China). Torchlight Hypersensitive ECL Western HRP Substrate(17046) was purchased from zen-bioscience (Chengdu, China). Goat Anti-Rabbit IgG (H+L) HRP was purchased from Affinity (Jiangsu, China). Antibodies against β-actin(AC026), Bax(A19684), Bcl2(A20777), Beclin1(A7353), LC3(A5618), mTOR(A11355), p-mTOR(AP0094), Akt(A18675), p-Akt(AP0980), PTEN(A19104), and HRP Goat Anti-Mouse IgG (H+L) (AS003) were supplied by the ABclonal. TRAP staining kit (CR2203125, Spec: 50T) and Secondary antibodies against Collagen II (GB23301) was purchased from Servicebio (Wuhan, China). DAB kit (ZLI-9018) was provided by ZSGB-BIO (Beijing, China). Antibody against Collagen II (NB600-844) was obtained from NOVUS (Shanghai, China). Rat IL-1β ELISA Kit(ZC-36391, Spec. 48 Test) and Rat MMP-13 ELISA Kit (ZC-36747, Spec. 48 Test) were purchased from Zhuocai Biotechnology (Shanghai, China). CCK-8 Kit (B3350A, 21169949, Spec. 5*100T) was provided by Biosharp (Anhui, China). Annexin V-APC/PI double staining apoptosis detection Kit (KGA1030) was obtained from KeyGEN (Jiangsu, China). The reference multi-element calibration standard and internal standard was purchased from National Center of Analysis and Testing for Nonferrous Metals and Electronic Materials (Beijing, China).

DHJSD: *Angelica pubescens* Maxim. (Duhuo, Chinese) 9g, *Asarum heterotropoides* F.Schmidt (Xixin, Chinese) 6g, *Saposhnikovia divaricata* (Turcz. ex Ledeb.) Schischk. (Fangfeng, Chinese) 6g, *Neolitse a cassia* (L.) Kosterm (Rougui, Chinese) 6g, *Gentiana macrophylla* Pall. (Qinjiao, Chinese) 6g, *Taxillus chinensis* (DC.) Danser (Sangjisheng, Chinese) 6g, *Eucommia ulmoides* Oliv. (Duzhong, Chinese) 6g, Cyathula officinalis K.C.Kuan, (Chuanniuxi, Chinese) 6g, *Paeonia lactiflora* Pall. (Baishao, Chin ese) 6g, *Rehmannia glutinosa* (Gaertn.) DC. (Dihuang, Chinese) 6g, *Angelica sinensis* (Oliv.) Diels (Danggui, Chinese) 6g, *Panax ginseng* C.A.Mey. (Renshen, Chinses) 6g, *Conioselinum anthriscoides ‘Chuanxiong’* (Chuanxiong, Chinese) 6g, *Smilax glabra* Roxb. (Fuling, Chinese) 6g, *Glycyrrhiza glabra* L. (Gancao, Chinese) 6g. All the herbs in the DHJSD were obtained from pharmacy department of Hospital of Chengdu University of Traditional Chinese Medicine.

Three doses of DHJSD (7*93g, 13*93g, 20*93g) were soaked in 2.5L, 3.5L and 5.5L of water for 60 minutes respectively, and then boiled. After boiling, each solution was concentrated to 1085ml, 1007.5ml, and 980ml, respectively, at a temperature of 50°C. Finally, low dose (0.6g/ml), medium d ose (1.2g/ml), and high dose (1.9g/ml) of DHJSD were obtained. Then, three different dosages of DHJSD were packaged in sterile bottles and stored at 4°C for future use.

### 2.2 Preparation of DHJSD-serum

This study was approved by the Experimental Animal Ethics Committee of Hospital of Chengdu University of Traditional Chinese Medicine (No.2023DL-005).Twenty adult male wistar rats (198g-220 g) were obtained from Chengdu Dossy Experimental Animals Co. LTD (Sichuan, China). Rats were bred for 1 week in an environment of 24±2°C, with free access to food and water, and a 12-hour cycle of light/darkness was maintained. Rats were randomly divided into two groups: group A and group B, with 10 rats in each group. The group A received high dose of DHJSD 2 times daily (1.5mL/100g·d) for 1 week. The same protocol was used with the group B, but the DHJSD was replaced with normal saline. One hour after the last administration, blood was collected from the abdominal aorta, and serum was separated by centrifugation, inactivated in a water bath at 56° C for 30 minutes, packaged, and frozen at -80 ° C until use. Then, all rats were euthanized in accordance with the ethical guidelines for animal welfare. See **Fig. 1**.

**Fig. 1.**
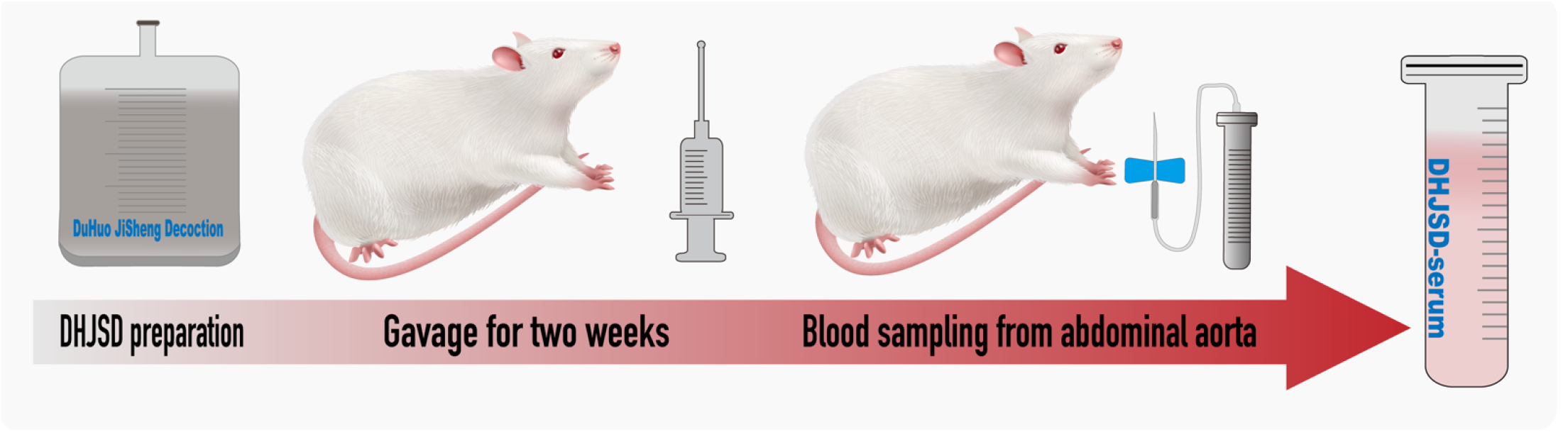
Schematic diagram of the preparation process of DHJSD-serum.

### 2.3 Chemical components and quality control of DHJSD

DHJSD-serum (Ds) components were detected using an LC-MS/MS system, ultra-high performance liquid chromatograph (Nexera UHPLC LC-30A, SHIMADZU, Japan) with mass spectrometer (TripleTOF5600+, AB SCIEX™). The liquid chromatography conditions were as follows: ChromCore 120 C18 Column (1.8μm 2.1mm*150mm), column temperature of 40°C, mobile phase A of 0.1% formic acid and mobile phase B of 100% ACN, flow rate of 0.3mL/min, and analysis time of 21min. Detection was performed using electrospray ionization (ESI) in negative ion mode. The ESI source conditions were as follows: Ion Source Gas1 (Gas 1): 50, Ion Source Gas2 (Gas 2): 50, Curtain Gas (CUR): 25, temperature of 450°C (negative ion), voltage of 4400V (negative ion), TOF MS scan range of 100-1200Da, product ion scan range of 50-1000Da, TOF MS scan accumulation time of 0.2s, product ion scan accumulation time of 0.01s. The second-level mass spectrometry was obtained using information-dependent acquisition (IDA) in high sensitivity mode, with a declustering potential (DP) of ±60V and a collision energy of 35±15eV.

### 2.4 PTEN siRNA

Chondrocytes were transfected with a specific PTEN siRNA synthesized by RiboBio(Guangzhou, China). A 1.5 mL centrifuge tube was prepared, and 15 μL siRNA was diluted with 360μL 1X riboFECTTMCP Buffer and gently mixed. Then 36μL riboFECTTMCP Reagent was added, gently blown and mixed, and incubated at room temperature for 15 min to prepare riboFECTTMCP transfection complex with a final siRNA concentration of 50 nmol for later use. The chondrocytes were seeded in a six-well plate. After the cells adhered to the wall, the supernatant in the wells was removed by suction, and the riboFECTTMCP transfection complex was added to an appropriate amount of complete medium without double antibody, mixed, and added to the well plate, and then incubated at 37°C and 5%CO_2_ for further use.

### 2.5 Cell treatment and CCK-8 assay

Cell grouping and treatments were shown in **Fig. 2**. CCK-8 assay was used to determine the optimal concentration in four different dosages of Ds (0, 5%, 10%, 15%, 20%). CCK-8 kit was used to detect the cell proliferation rate, and the operation was strictly in accordance with the instructions. Rat primary chondrocytes(Rat-icell-s003, icell) (5×10^4^/mL, 100μL/ well) in each group (4 wells in each group) were seeded in 96-well plates (edge wells were filled with sterile PBS) and cultured for 48 hours at 37°C with 5% CO_2_. After 48h of treatment, the supernatant was aspirated and discarded. CCK-8 reagent was diluted 1:10 in serum-free medium, and 110μL diluted CCK-8 solution/well was added. The culture plate was gently shaken several times, and the culture was continued for 2 hours at 37°C and 5% CO_2_. The absorbance values of each well were measured at a wavelength of 450nm using a microplate reader.

**Fig. 2.**
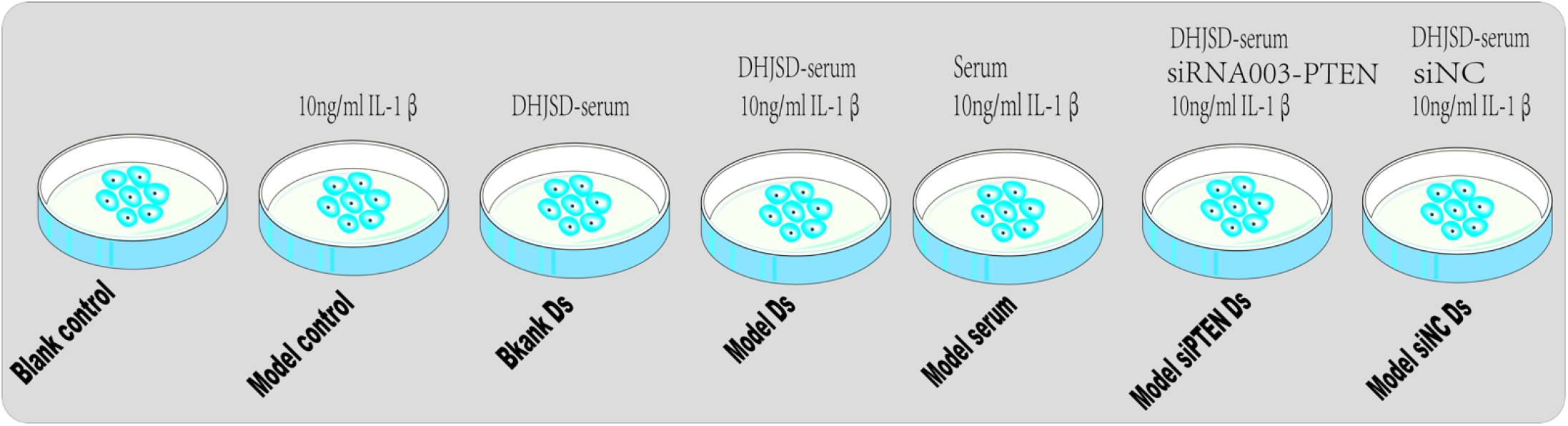
Schematic representation of cell grouping and pretreatment.

### 2.6 Measurement of cell apoptosis

The cells of each group were collected. After the cells were resuspended in 500μL Binding Buffer, 5μL Annexin V-APC/PI was added to blow gently, and then 5 μl PI was added to mix. The reaction was carried out at room temperature in the dark for 15min, and then detected and analyzed.

### 2.7 Determination of intracellular zinc levels

Sample was placed in the Teflon vessel according to the grouping. Concentrated nitric acid (5 mL) was added. Perform digestion according to the standard operation steps of microwave digestion machine (Shanghai, China). The ICP-MS/MS (iCAP TQ, ThermoFisher, USA) was applied to determine the level of zinc in chondrocytes. The tuning fluid was used to adjust and optimize the instrument parameters before each experiment to make the sensitivity, resolution and stability meet the requirements, and CCT(He/O_2_ mode) was used for analysis. The main working parameters and acquisition conditions of the instrument are as follows ^[23, 24]^: Tuning mode: STD/KED; Dwell time: 0.1 s; Peristaltic Pump Speed: 40 rpm; Sample introduction time: 40 s ; Plasma Power: 1550 W; Sampling Depth: 5.0 mm; Nebulizer Flow:0.98 L/min; Cool Flow:14.0 L/min; Auxilliary Flow:0.8 L/min; Spray Chamber Temperature: 2.7 °C; Torch Horizontal Position: 0.16 mm; Torch Vertical Position: -0.53 mm; He flow: 4.55 mL/min; O_2_flow: 0.3125 mL/min; D1 Lens: -350 V; D2 Lens: - 350 V; Repeat times: 3 times.

### 2.8 Animal procedure

#### 2.8.1 Establishment of OA rat model

This study was approved by the Experimental Animal Ethics Committee of Hospital of Chengdu University of Traditional Chinese Medicine (No.2023DL-005). A total of 25 adult male wistar rats (198g-220 g) were obtained from Dossy (Chengdu, Sichuan, China). Rats were bred for 1 week in an environment of 24±2°C, with free access to food and water, and a 12-hour cycle of light/darkness was maintained. Rats were randomly divided into the sham control group (where only the joint cavity was exposed; n= 5), model control group (n = 5), model HD(high dose) group (n = 5), model MD(medium dose) group (n = 5), model LD(low dose) group (n = 5).

The rat OA model was established by the Hulth method under anesthesia of pentobarbital sodium. As showed in **Fig. 3**, the incision was made from the medial side of right posterior knee joint of the rats after anesthetized intraperitoneally with pentobarbital. The muscles as well as ligaments were separated, and the joint cavity was exposed. The medial collateral ligament, anterior and posterior cruciate ligament, as well as the medial meniscus was were cut off. Then, the incision was sutured. After operation, each rat was intramuscular injection with penicillin (400 000 U/d) for 3 consecutive days, and then kept in the cage.

**Fig. 3.**
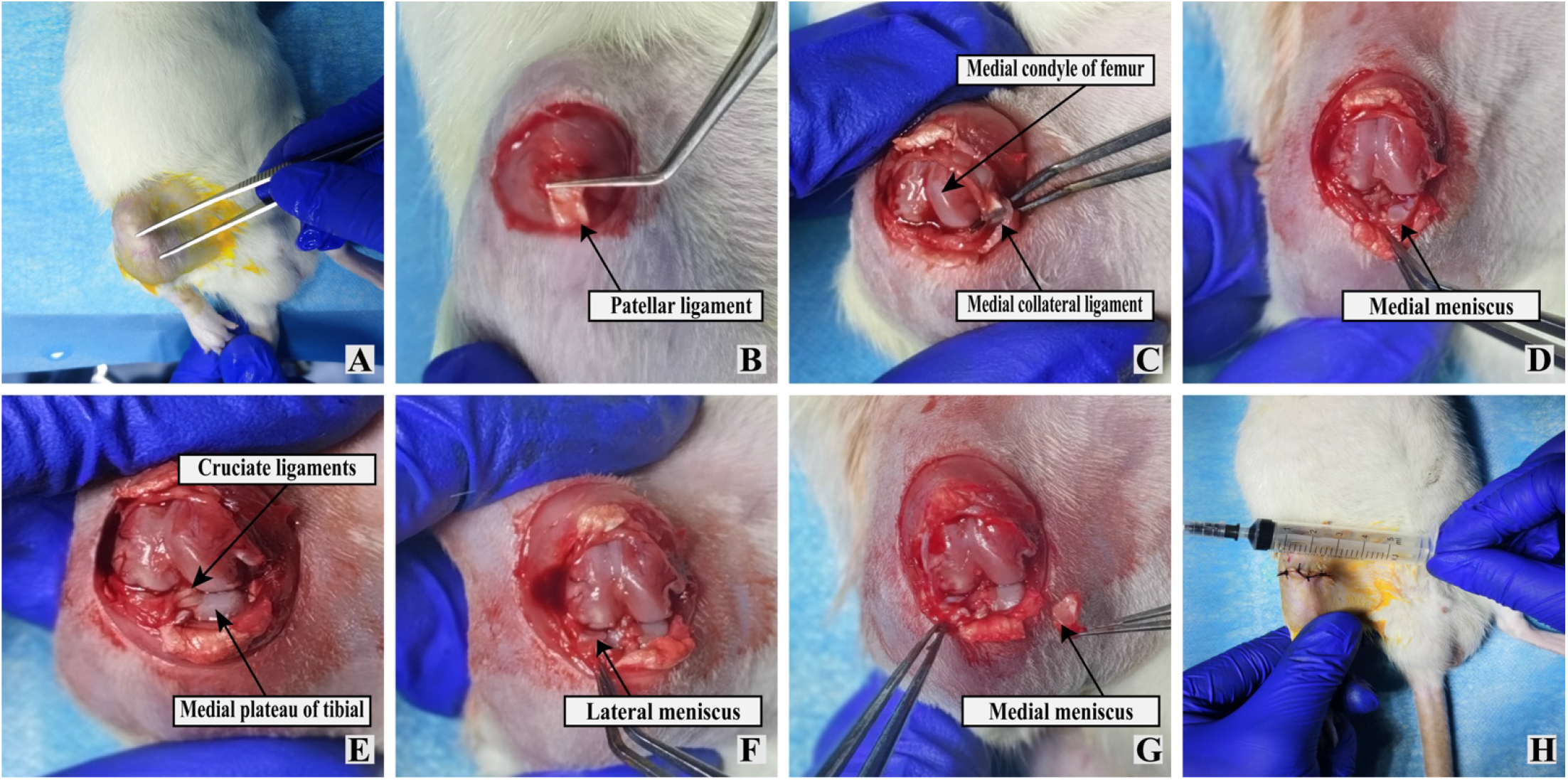
Procedure for establishing a rat KOA model using the Hulth method. (A) Incision site; (B) Exposure of patellar ligament; (C) Cutting of patellar ligament and exposure of medial collateral ligament; (D) Cutting of medial collateral ligament and exposure of medial meniscus; (E). Removal of medial meniscus and exposure of cruciate ligament; (F) Cutting of cruciate ligament and retention of lateral meniscus; (G) Medial collateral ligament and cruciate ligament have been cut, medial meniscus has been removed, and lateral meniscus is fully retained; (H) Suturing of incision.

#### 2.8.2 Animal treatments

The model HD group, model MD group and model LD group were given with the same volume of DHJSD at high, medium and low dose by gavage 2 times daily (1.5mL/100g·d) for 4 weeks from the first day after operation, respectively (**Fig 4**). The same protocol was used with the sham group and the model control group while the DHJSD was replaced with normal saline. After treatment, all rats were euthanized in accordance with the ethical guidelines for animal welfare. The specimens of blood and knee joints were obtained for further tested.

**Fig. 4.**
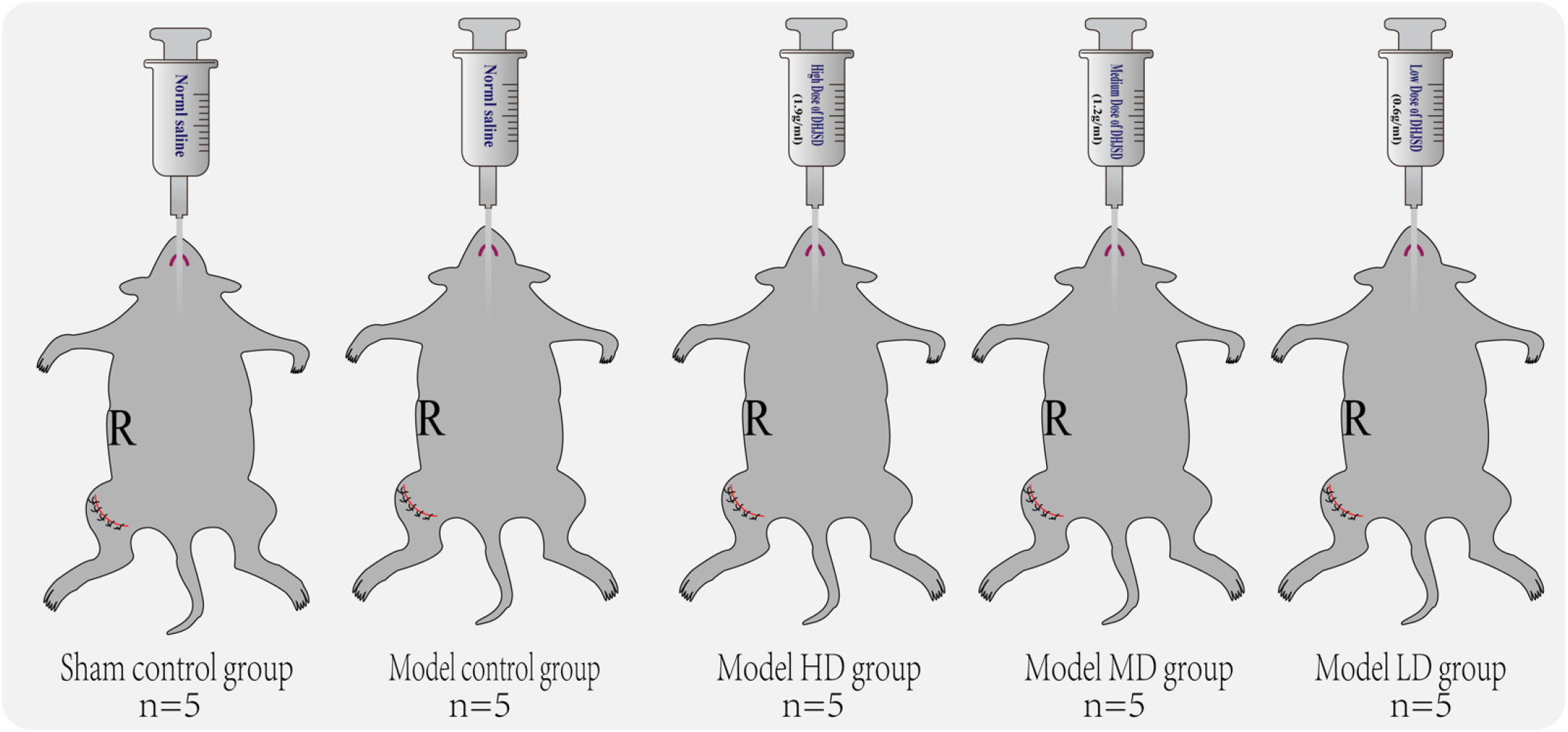
Schematic representation of animal grouping and treatment.

### 2.9 Histological analysis and macroscopic observation

Hematoxylin-eosin staining, masson staining, immunohistochemical staining and trap staining were used to observe the cartilage morphology under light microscope. The autophagosomes of chondrocytes were observed by transmission electronic microscopy.

### 2.10 Elisa

Quantification of the levels of MMP-13 and IL-1β in each group of serum were performed using the corresponding enzyme-linked immunosorbent assay (ELISA) kit. The absorbance at 450 nm was measured by a microplate reader (SpectraMAX Plus384, Molecular Devices, USA).

### 2.11 Real-time PCR

Total cellular RNA was isolated using animal total RNA isolation kit (RE-03014, Foregene, China) according to the manufacture’s instruction. Complementary DNA (cDNA) was reverse-transcribed with PrimeScript-RT reagent kit (RR047A, Takara, Japan).The mRNA levels of Beclin-1, Bax, Bcl-2 and PTEN were detected by RT-PCR with TB Green TM Premix Ex TaqTM II(Tli RNaseH Plus) (RR820A, Takara, Japan). The complete gene sequences showed in **Table 2** were searched from the National Center for Biotechnology Information (NCBI) database, and the specific primers were designed and screened by Primer Premier software. All primers were designed and synthesized by Sangon Bioengineering Technology(Shanghai, China), and purified by ULTRAPAGE. Data were analyzed by 2^−ΔΔCT^ method.

**Table 2.**
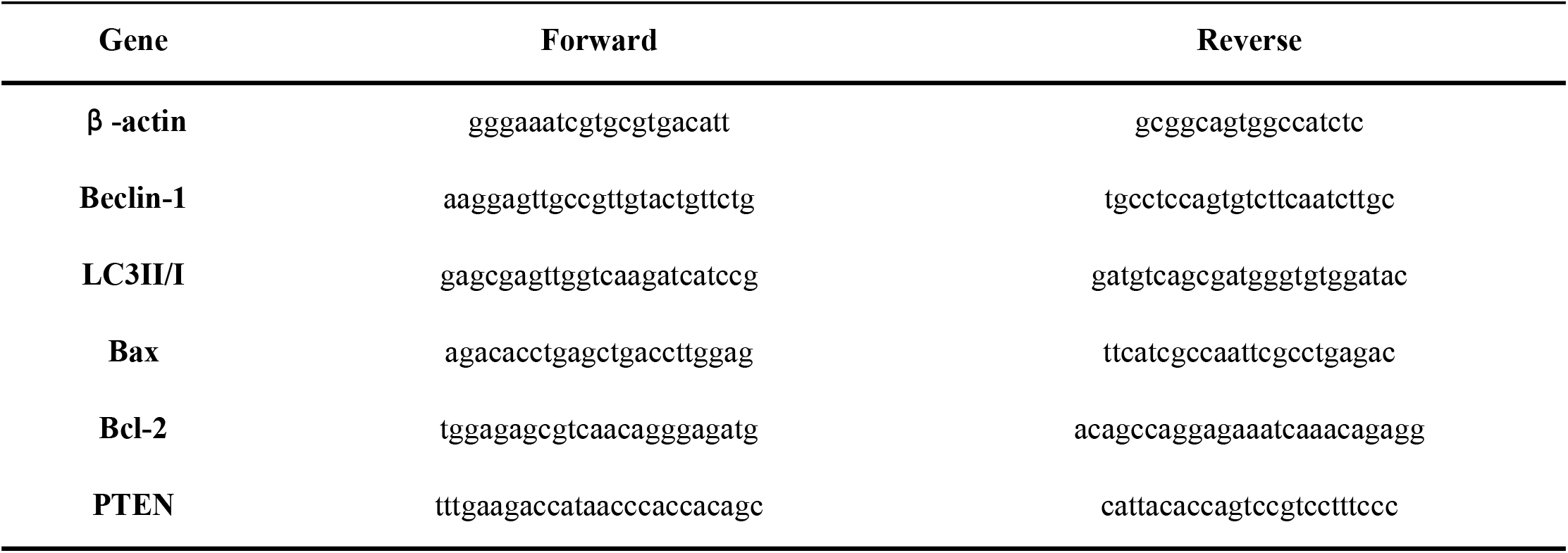
Primer sequence.

### 2.12 Western Blot

The protein levels of PTEN, Akt, p-Akt, mTOR, p-mTOR, Bax, Bcl-2, Beclin-1 and LC3-II/I in chondrocytes were detected by Western blot. Total cellular proteins were extracted using RIPA lysates. The proteins were transferred to PVDF membrane by wet rotation method and blocked at room temperature for 1 h. The primary antibody was incubated overnight and the membrane was washed 3 times with TBST for 5 min each time. The secondary antibody was incubated at room temperature for 2 h, and the membrane was washed 3 times with TBST for 10 min each time. Then the ECL developer solution was uniformly added to the membrane for exposure development. The bands were scanned by exposure using Tianneng GIS chassis control software V2.0, and the results were expressed as the relative expression of the target protein.

### 2.13 Statistical analyses

SPSS 26 (IBM® SPSS® Statistics) was adopted to analyze the data generated in the charts in this experiment, and figures were made by using GraphPad Prism v9.5.1 (GraphPad Software, Inc.). All data are presented as the mean ± SEM, and all experiments were performed three times. Differences among seven groups were analyzed by one-way analysis of variance followed by Fisher’s post hoc test. Differences between two groups were statistically analyzed using an unpaired, two-tailed Student’s t-test. P<0.05 was considered to indicate a statistically significant difference.

## 3. Result

### 3.1 Qualitative analysis of compounds contained in DHJSD-serum

A total of 173 compounds were detected. Substances with a total score of 100, including formula, ontology, reference and RT, were listed as in **Table 3**.

**Table 3.** The compounds contained in DHJSD-serum with a total score of 100.

| Formula | Ontology | Reference (m/z) | RT (min) | Total score |
| --- | --- | --- | --- | --- |
| C<sub>17</sub>H<sub>22</sub>O<sub>11</sub> | Iridoid glycosides | 401.10849 | 6.1149 | 100 |
| C<sub>21</sub>H<sub>44</sub>NO<sub>6</sub>p | Glycerophosphoethanolamines | 436.28336 | 15.86822 | 100 |
| C<sub>20</sub>H<sub>36</sub>N<sub>4</sub>O<sub>8</sub> | N-acyl amines | 461.26059 | 17.3248 | 100 |
| C<sub>9</sub>H<sub>17</sub>NO<sub>5</sub> | Fatty acids | 218.1032 | 1.7611 | 100 |
| C<sub>6</sub>H<sub>12</sub>O<sub>6</sub> | Monosaccharides | 179.056 | 1.239233 | 100 |
| C<sub>5</sub>H<sub>14</sub>NO | Cholines | 104.10699 | 19.25602 | 100 |
| C<sub>8</sub>H<sub>11</sub>NO<sub>3</sub> | Pyridoxines | 170.05896 | 5.367417 | 100 |
| C<sub>15</sub>H<sub>22</sub>O<sub>3</sub> | Tertiary alcohols | 249.08504 | 6.836583 | 100 |
| C<sub>33</sub>H<sub>40</sub>N<sub>2</sub>O<sub>9</sub> | Yohimbine alkaloids | 609.33337 | 16.61178 | 100 |
| C<sub>5</sub>H<sub>4</sub>N<sub>4</sub> | Purines and purine derivatives | 119.036 | 1.36025 | 100 |
| C<sub>9</sub>H<sub>10</sub>O<sub>3</sub> | Alkyl-phenylketones | 165.05571 | 7.313766 | 100 |

### 3.2 Effect of DHJSD on the viability of chondrocytes incubated with or without IL-1β

The cytotoxicity of Ds (0, 5%, 10%, 15% and 20%) on rat chondrocytes was determined for 48 hours. The results showed that the cell viability of Ds (10 %) treatment for 48 hours was stronger than that of the other four groups. Chondrocytes viability rate was inhibited by IL-1β (10ng/ml), and this effect was reversed when pretreatment with Ds (10%). However, this effect was significantly attenuated after PTEN gene silencing(**Fig. 5**). Therefore, Ds (10%) was used for subsequent experimental studies.

**Fig. 5.**
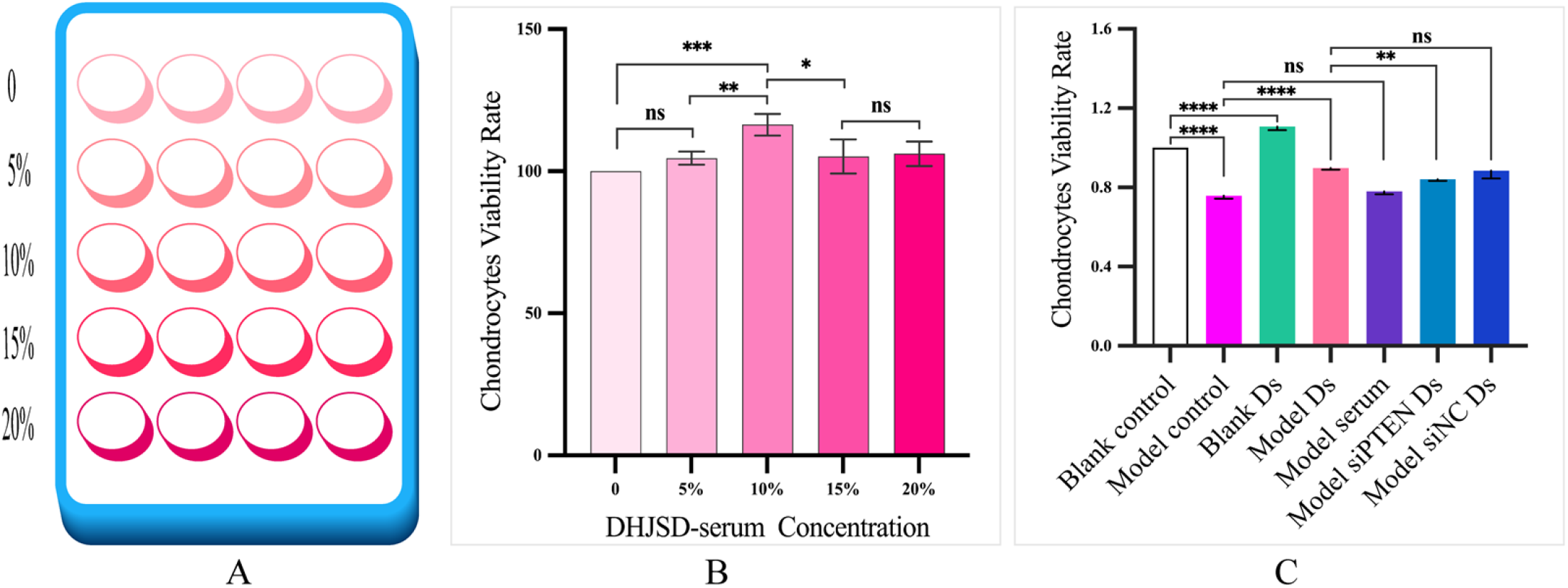
CCK8 was used to screen the optimal concentration (A, B) and the effect of Ds (10%) on the cell viability of each group (C). Ds=DHJSD-serum. (n=3, mean±SD; *p<0.05, **p<0.01, ***p<0.001, ****p<0.0001,^ns^p>0.05).

### 3.3 Effect of DHJSD on apoptosis of chondrocytes *in vivo and vitro*

In our study, the effect of IL-1β treatment on chondrocytes apoptosis was further confirmed. Ds significantly reduced the level of apoptosis, while this effect was significantly depressed after PTEN gene silencing (**Fig. 6**). In addition, Bax and Bcl-2 protein expressions were significantly up-regulated after IL-1β treatment, but the effect was reversed when pretreatment with Ds. Synthesis of specific siRNA PTEN can silence PTEN expression and inhibit the anti-apoptotic effect of Ds (**Fig. 7A, 7C, 7D**).To further verify this phenomenon, experiments *in vivo* were performed. The results showed that DHJSD can inhibit the expression of Bax and Bcl-2 proteins in chondrocytes of rats with KOA induced by Hulth method, and there was a certain dose dependence (**Fig. 7B, 7E, 7F, 7G, 7H**). These results indicated that DHJSD produced a significant anti-apoptotic effect, which may be achieved by targeting PTEN.

**Fig. 6.**
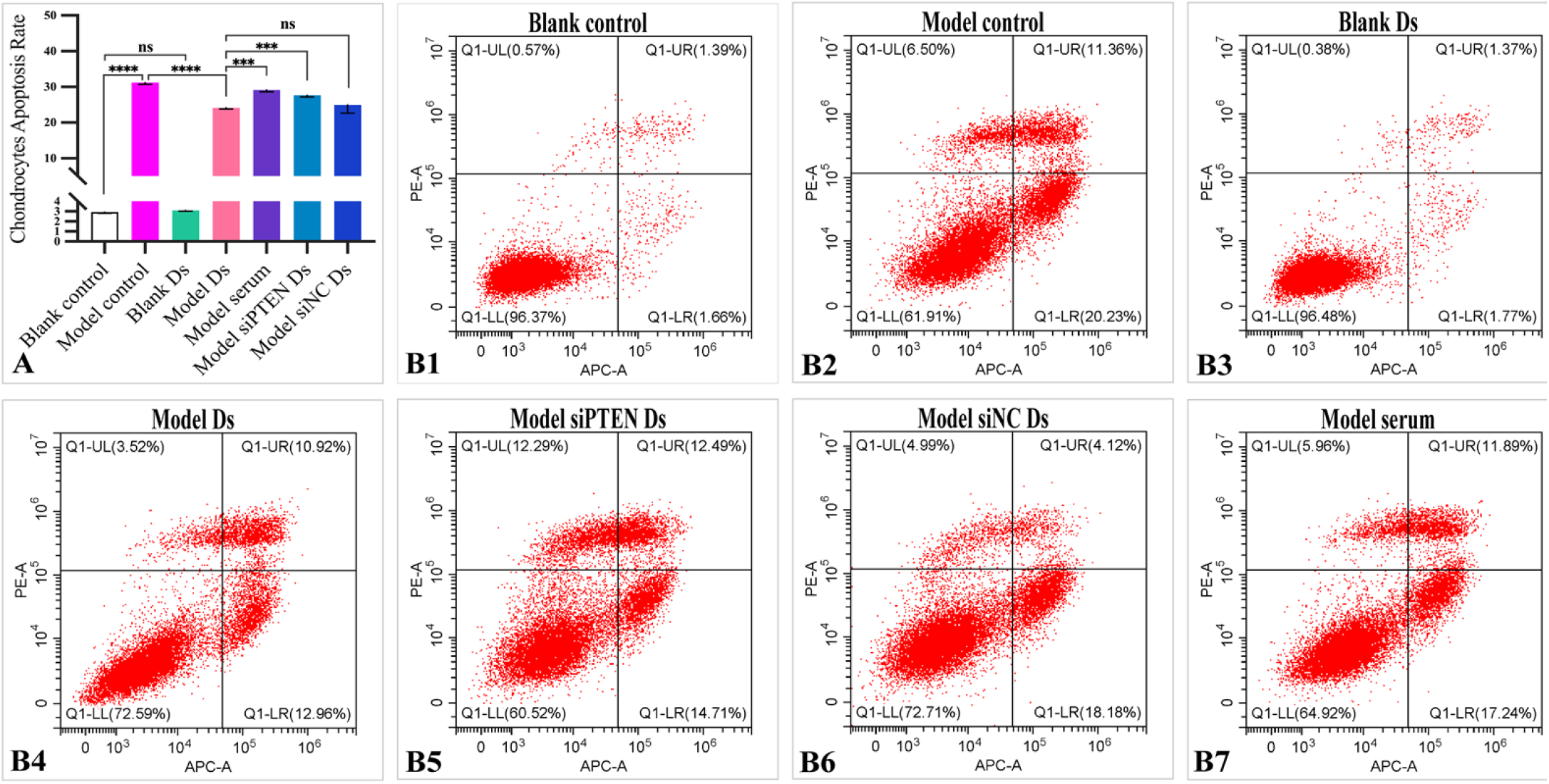
Effect of DHJSD on apoptosis of chondrocytes *in vitro*. (A) According to the grouping, the chondrocytes were pretreated with IL-1β, Ds and the riboFECT™CP transfection complex. After 48 hours of cultured, the chondrocytes apoptosis was determined by flow cytometry in each group. (B1-B7) Scatter plot of chondrocyte apoptosis. (n=3, mean±SD; ***p<0.001, ****p<0.0001,^ns^p>0.05).

**Fig. 7.**
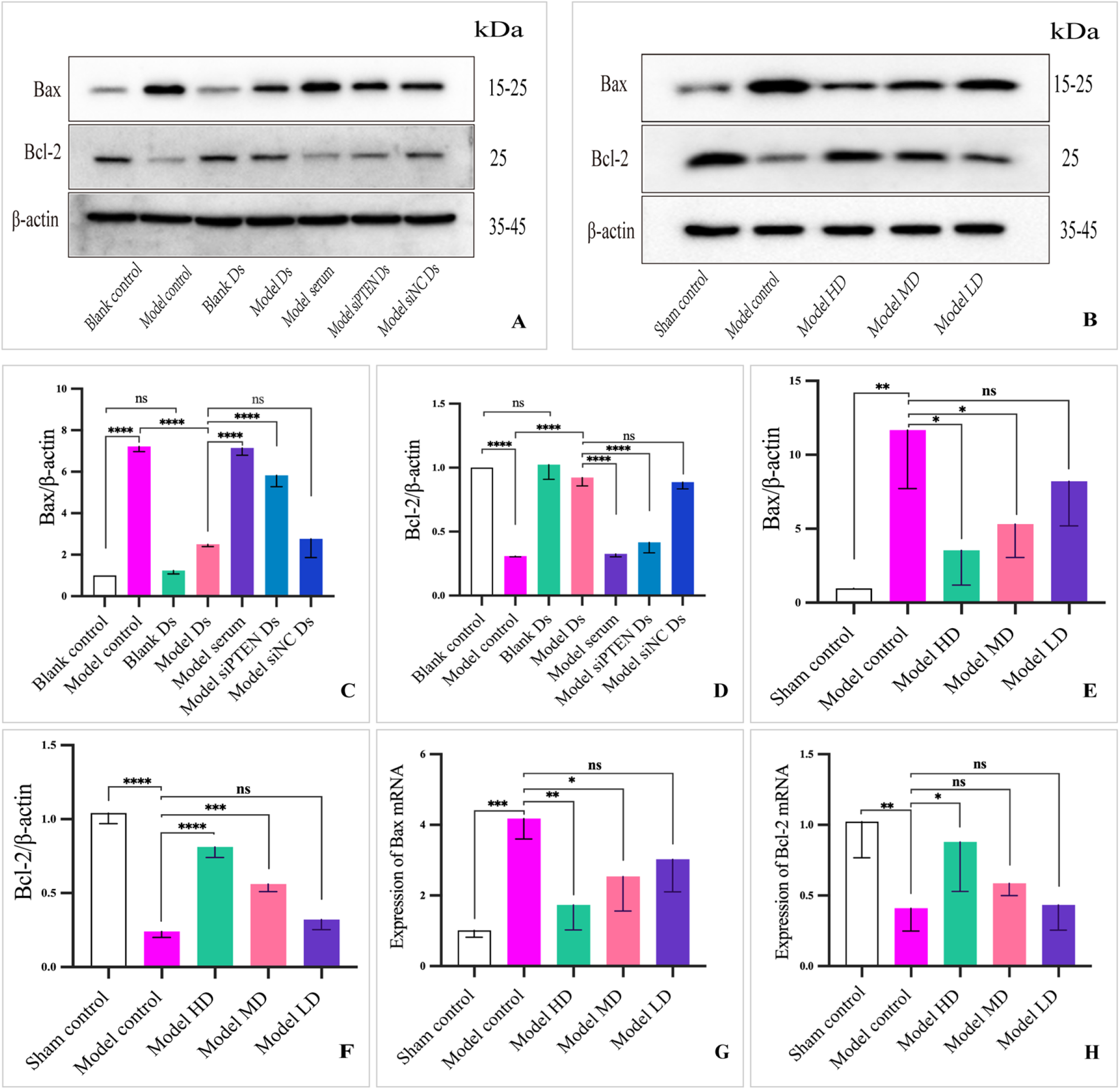
Effect of DHJSD on the expression of apoptotic proteins in chondrocytes. (A, C, and D) According to the grouping, the chondrocytes were pretreated with IL-1β, Ds and the riboFECT™CP transfection complex. After 48 hours of culture, the expressions of Bax and Bcl-2 in each group were detected. (B, E, F,G, H) After 4 weeks of pretreatment with different dosages of DHJSD in rats with KOA established by Hulth method, the expressions of Bax and Bcl-2 in knee chondrocytes were detected. (n=3, mean±SD; *p<0.05, **p<0.01, ***p<0.001, ****p<0.0001,^ns^p>0.05).

### 3.4 Effect of DHJSD on the level of autophagy in chondrocytes

As shown in **Fig. 8A-J**, the level of autophagy in chondrocytes was decreased after IL-1β treatment. Meanwhile, Ds significantly increased the level of autophagy in chondrocytes stimulated by IL-1β. However, after silencing PTEN gene, this effect was significantly weakened. Additionally, the protein expressions of Beclin-1 and LC3II/I were significantly down-regulated after IL-1β treatment, but these trends were reversed when pretreatment with Ds. Synthesis of specific siRNA PTEN can silence PTEN expression and inhibit Ds-induced autophagy. The results also showed that DHJSD could up-regulate the protein expression of Beclin-1 and LC3II/I in chondrocytes of rats with KOA induced by Hulth model in a dose-dependent manner *in vivo*. What’s more, transmission electron microscopy showed that after treatment with DHJSD, there was a significant increase in autophagic vesicles in the cytoplasm, which was positively correlated with the dose of DHJSD. These results suggested that DHJSD induced autophagy of chondrocytes, which may be mediated by the targeted regulation of PTEN. Collectedly, it was reasonable to speculate that DHJSD promoted autophagy by targeting PTEN according to the results.

**Fig. 8.**
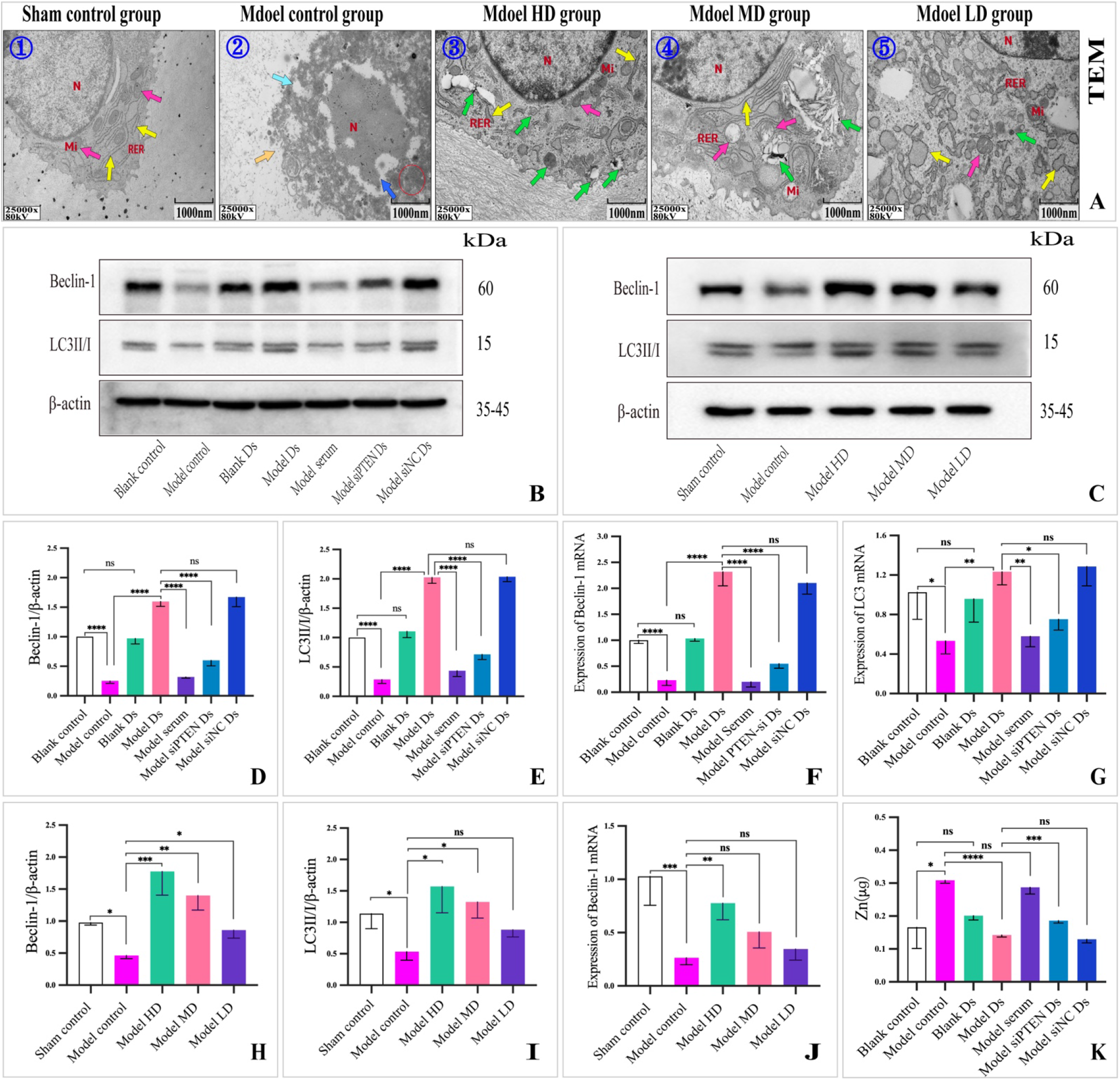
Effect of DHJSD on autophagy and level of zinc in chondrocytes. (A). After 4 weeks of pretreatment with different dosages of DHJSD in rats with KOA induced by Hulth method, the morphological and structural changes of cartilage organelle were observed by transmission electron microscope. Sham group: cell morphology, mitochondria morphology, rough endoplasmic reticulum morphology and structure were normal. In the model group, chondrocytes were necrotic, the cell body shrank and became smaller, the cell membrane was discontinuous, the cytoplasm was concentrated, the cytoplasmic content was lost in some areas, the nucleus was lysed, and the organelles were dissolved and blurred. In the Model HD group, the morphological structure of cells, the morphological structure of mitochondria were normal, the rough endoplasmic reticulum of some cells was dilated, and there were more autophagosomes in the cytoplasm. A-❶: 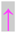 Normal mitochondria, 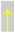 Normal endoplasmic reticulum; A-❷: 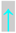 Loss of cytosolic contents,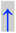 Breakdown of nuclear membrane,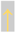 Discontinuous cell membrane,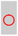 Cytoplasmic pyknosis; A-❸: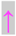 Mitochondria, 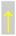 Endoplasmic reticulum, 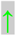 Autophagosomes; A-❹: 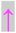 Mitochondria with swollen morphology, 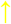 Endoplasmic reticulum, 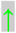 Autophagosomes; A-❺: 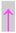 Mitochondria, 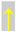 Endoplasmic reticulum with distend morphology, 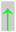 Autophagosomes. (B, D, E, F, G) Pretreatment with IL-1β, Ds and the riboFECT™CP transfection complex according to groups, Western blot and RT-PCR were used to detect Beclin-1 and LC3 in cartilage of each group after 48 hours of culture. (C, H, I, J) After 4 weeks of pretreatment with different dosages of DHJSD in rats with KOA induced by Hulth method, the expressions of Beclin-1 and LC3 in chondrocytes were determined. (n=3, mean±SD; *p<0.05, **p<0.01, ***p<0.001, ****p<0.0001,^ns^p>0.05).

### 3.5 Effect of DHJSD on zinc homeostasis in chondrocytes

A study showed^[4]^ that DHJSD can down-regulate the level of zinc in chondrocytes and inhibit the expression of MMP-13 and other proteins. To verify the effect of DHJSD on intracellular zinc level, ICP-MS/MS was used to measure the level of zinc in chondrocytes in this study. The results showed that the level of zinc in chondrocytes increased after IL-1β treatment. However, Ds reduced intracellular zinc level significantly, and this effect was significantly receded after PTEN gene silencing (**Fig. 8K**).

### 3.6 Effect of DHJSD on PTEN/Akt/mTOR signaling pathway

It is well known that the Akt/mTOR signaling axis is closely related to autophagy, and PTEN has a direct regulatory effect on this signaling pathway. In the present study, we found that the pretreatment of Ds promoted PTEN expression and reduced Akt and mTOR phosphorylation in IL-1β-stimulated chondrocytes. However, when PTEN gene was silenced, the inhibitory effect of Ds on Akt and mTOR phosphorylation was weakened. It is worth mentioning that Ds has no significant effect on PTEN gene, Akt and mTOR phosphorylation levels in normal chondrocytes. Also, the results were confirmed in vivo. See **Fig. 9**.

**Fig. 9.**
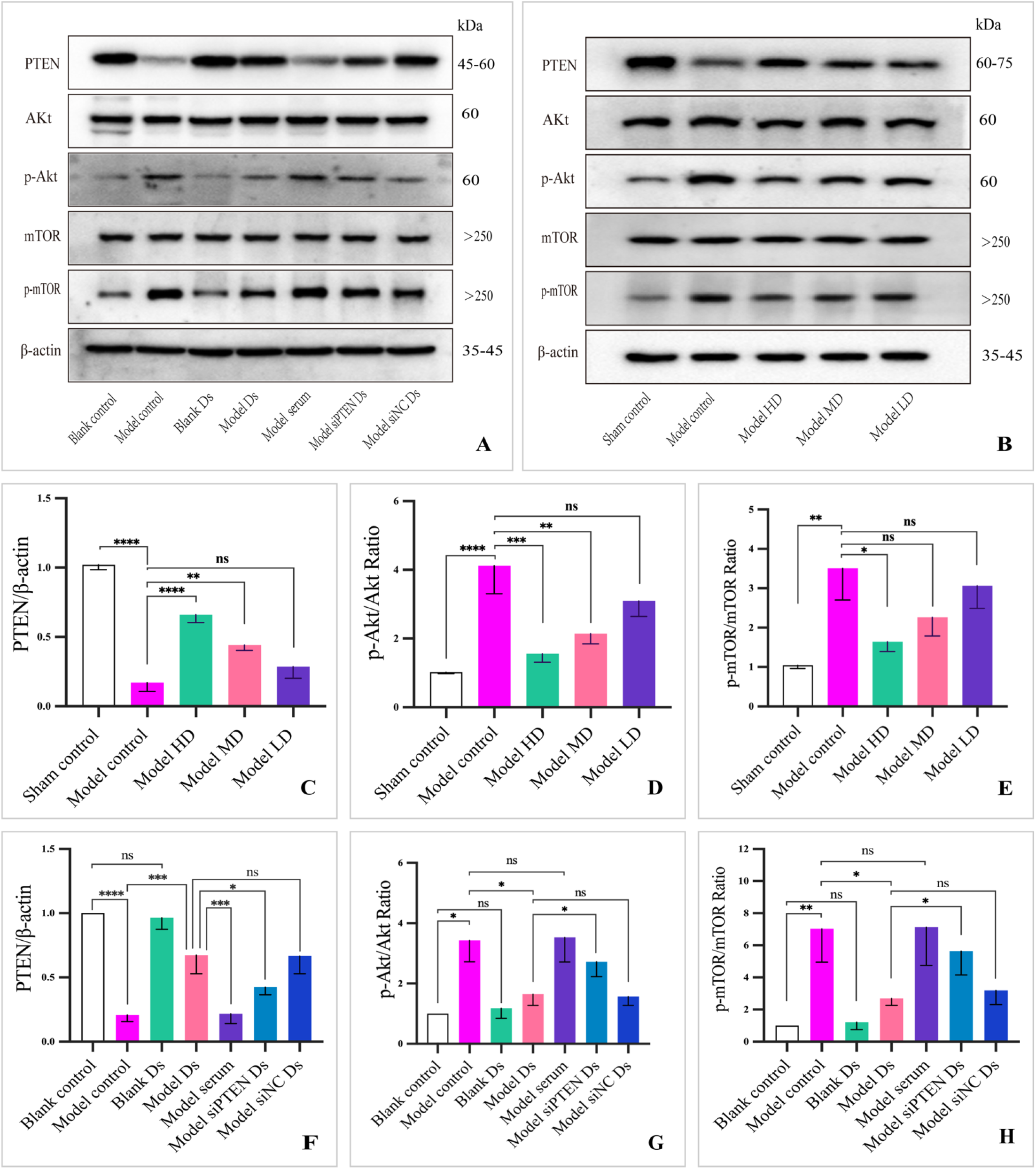
Effect of DHJSD on PTEN/Akt/mTOR signaling pathway in chondrocytes *in vitro and vivo*. (A, F, G, H) According to the grouping, the chondrocytes were pretreated with IL-1β, Ds and the riboFECT™CP transfection complex. After 48 hours of culture, the expressions of PTEN, p-Akt, Akt, p-mTOR and mTOR in each group were detected. (B, C, D, E) After 4 weeks of pretreatment with different dosages of DHJSD in rats with KOA induced by Hulth method, the expressions of PTEN, p-Akt, Akt, p-mTOR and mTOR in knee chondrocytes were detected. (n=3, mean±SD; *p<0.05, **p<0.01, ***p<0.001, ****p<0.0001,^ns^p>0.05).

### 3.7 DHJSD alleviated cartilage degeneration in OA rat

To observe the chondroprotective effects of DHJSD *in vivo*, a rat OA model was established by Hulth method. As shown in **Fig. 10**, after 4 weeks of Hulth surgery, the model group exhibited severe loss of type II collagen, cartilage destruction, osteoclast formation, and collagen fiber formation, and had a higher Mankin score compared to the sham control group. However, these changes were mild after pre-treatment with a high dose of DHJSD. The results indicated that DHJSD had a certain inhibitory effect on cartilage destruction.

**Fig. 10.**
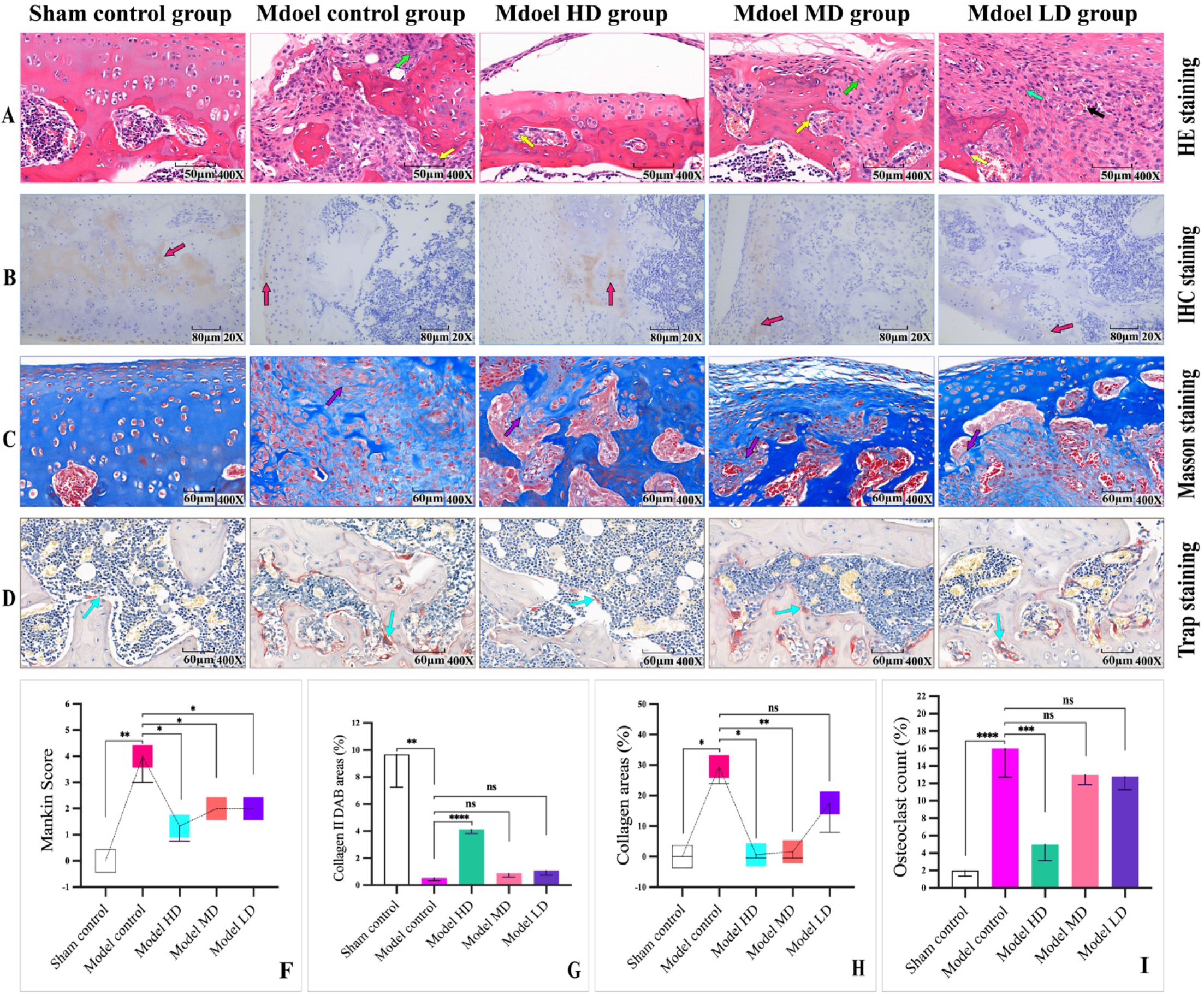
DHJSD alleviated cartilage degeneration in rats with KOA. After 4 weeks of pretreatment with different concentrations of DHJSD in rats with KOA induced by Hulth method, the morphological and structural changes of cartilage tissue were observed and analyzed. (A) HE staining, 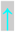 Osteoclasts, 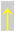 fibroblasts, 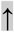 new capillaries; (B) collagen II IHC staining, 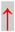 collagen II; (C) Masson staining, 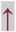 collagenous fiber; (D) Trap staining, 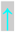 Osteoclasts. In the model group, the joint tissue surface exhibited cartilage erosion, with complete loss of the local cartilage layer structure, a high number of bone cell necrosis, reduced type II collagen levels, increased collagen fiber tissue proliferation deposition area, and a higher number of osteoclasts. After pretreatment with a high dose of DHJSD, a mild cartilage erosion was observed on the joint tissue surface (Fig. 9F, **p<0.01 vs Model control), with a small number of cartilage cell necrosis. There was a significant increase in the expression of type II collagen (Fig. 9G, *p<0.05 vs Model control), and a significant reduction in the deposition area of collagen fiber tissue and the number of surrounding osteoclasts compared to the model group (Fig. 9H, *p<0.05, Fig. 9I, ***p<0.001), as demonstrated by the results of this study. (n=3, mean±SD; *p<0.05, **p<0.01, ***p<0.001, ****p<0.0001,^ns^p>0.05).

### 3.8 Effects of DHJSD on MMP13 and IL-1β in rats with KOA

Besides, we investigated the effect of DHJSD on inflammatory factors in rats with KOA induced by Hulth method. As shown in the **Fig. 11**, the levels of MMP-13 and IL-1β in the serum, as well as the expression area of MMP-13 in the cartilage matrix were increased in four weeks after the Hulth procedure. However, the corresponding indicators showed varying degrees of reduction in rats with KOA pretreated with DHJSD, and the differences were statistically significant. These results further confirm that DHJSD can reduce the expression of MMP-13 in the cartilage matrix and serum of rats with KOA.

**Fig. 11.**
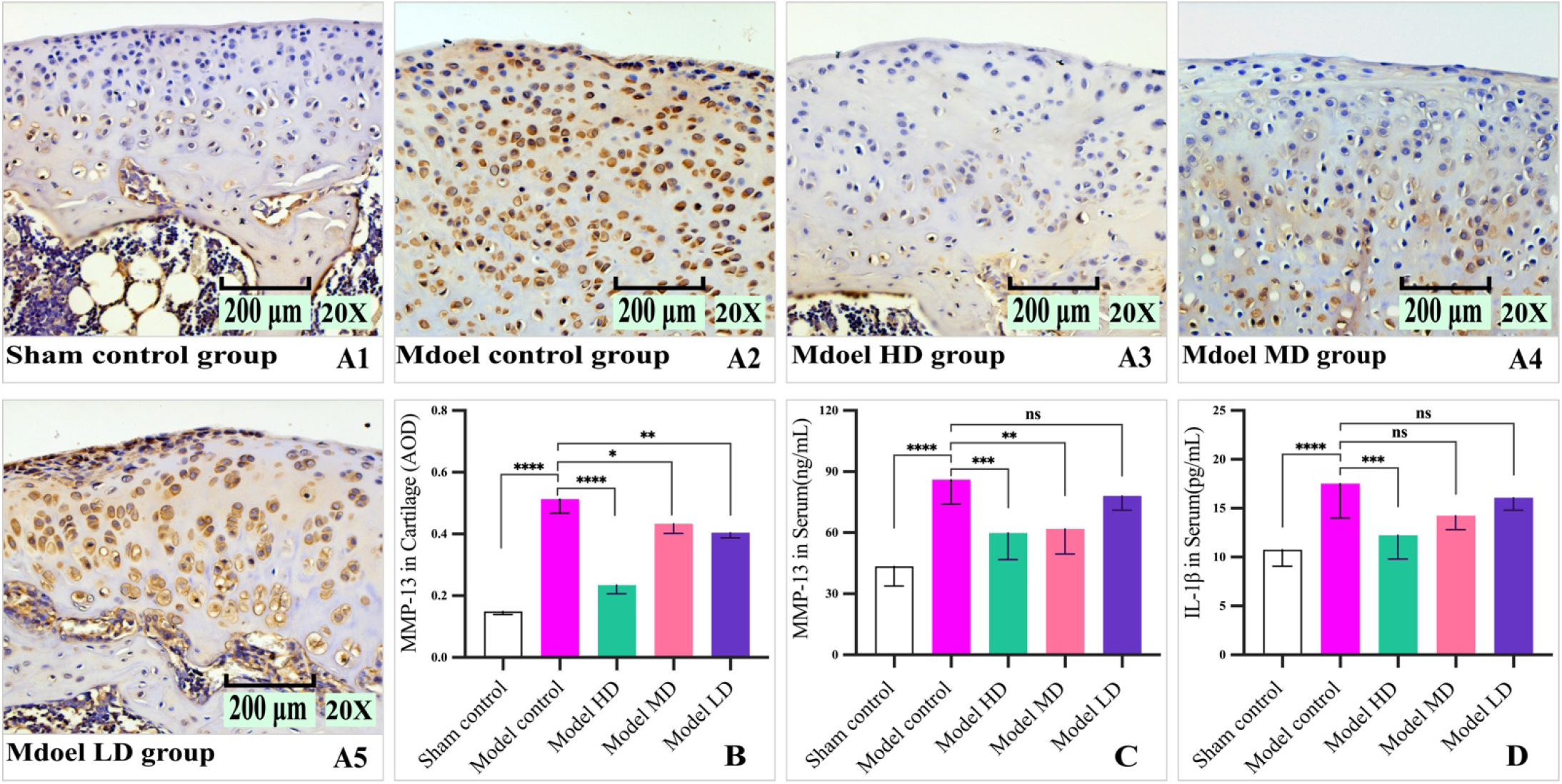
DHJSD decreased the levels of MMP-13 and IL-1β in OA rat. After 4 weeks of pretreatment with different dosages of DHJSD in rats with KOA established by Hulth method, the levels of MMP-13 and IL-1β in the serum, as well as the expression area of MMP-13 in the cartilage matrix were observed and analyzed. (A1-A5) Histological images of MMP-13 immunostaining in the cartilage matrix. (B) The statistical results of MMP-13 immunostaining in the cartilage matrix. (C, D) The statistical results of MMP-13 and IL-1β levels in serum measured by Elisa, respectively. The model control group: exhibited increased expression of MMP-13 in the cartilage matrix, and elevated levels of MMP-13 and IL-1β in serum. All three different dosages of DHJSD could inhibit the expression of MMP-13 in the cartilage matrix (B). However, only the high-dose group showed a simultaneous decrease in the levels of MMP-13 and IL-1β in serum (C, D: ***p<0.001 vs Model control). (n=3, mean±SD; *p<0.05, **p<0.01, ***p<0.001, ****p<0.0001,^ns^p>0.05).

## 4. Discussion

Inflammatory cytokine cascade activation, chondrocyte apoptosis, imbalance of subchondral bone remodeling, and extracellular matrix degradation are the four major pathological and etiological characteristics of KOA ^[25]^. Subchondral bone remodeling, delaying chondrocyte apoptosis, and inhibiting matrix degradation are the main research directions for studying cartilage degeneration. Currently, non-steroidal anti-inflammatory drugs (NSAIDs) are the most widely used and effective drugs in the conventional clinical treatment of KOA^[26, 27]^. However, the long-term use of these drugs has side effects such as gastrointestinal ulcers and cardiovascular adverse events^[28]^, which are concerning. Specific treatments for osteoarthritis, such as glucosamine, hyaluronic acid, and platelet-rich plasma injections, are controversial and either have conflicting guidelines or are not recommended owing to lacking in evidence-based medicine for the long-term efficacy^[29]^. Therefore, in the current situation, the use of natural methods, including traditional Chinese medicine formulas, is one of the important directions for the prevention and treatment of KOA. The DHJSD has a long history of use in treating KOA, but its mechanism of action has not been fully elucidated.

Degenerative factors run through the whole process of OA, and soft tissue mechanical abnormality is one of the initiating factors of OA. Therefore, the classic Hulth method was used to establish the rat KOA model in this study. IL-1β plays a key role in OA pathogenesis, and it is a well-established method for IL-1β to be used to establish an in vitro cellular OA model. In the present study, both the Hulth-established rat KOA model *in vivo* and IL-1β-induced cell model significantly replicated human OA-like features, and the Hulth-established rat OA-like features were attenuated by DHJSD.

This study aims to investigate the mechanism of DHJSD in the prevention and treatment of KOA in IL-1β-induced chondrocytes and rats with KOA established by Hulth method. The results showed that DHJSD could inhibit IL-1β and MMP-13, which was consistent with the results of previous studies, and confirmed the anti-inflammatory effect of DHJSD. This study in vivo also found that DHJSD could alleviate the destruction of articular cartilage layer, repressed the formation of osteoclasts, reduce the deposition of collagen fibers, and prevent the degradation of type II collagen in articular cartilage tissue. Moreover, DHJSD could up-regulate the expression of PTEN protein and mRNA, inhibit Akt/mTOR signaling pathway, promote the expression of LC3 and Belcin-1 in chondrocytes, inhibit the expression of Bax and Bcl-2 in chondrocytes, and reduce the level of zinc in the chondrocytes.

Autophagy is a key important protective mechanism for cells to avoid their own apoptosis^[30, 31]^. Moreover, the PTEN/Akt signaling axis is closely related to autophagy^[11, 17, 32]^. After IL-1β treatment, the expression of PTEN, LC3 and Beclin-1 was down-regulated, and the phosphorylation of Akt and mTOR was increased. Pretreatment of IL-1β-stimulated chondrocytes with Ds increased the expression of PTEN, LC3 and Beclin-1 and decreased the phosphorylation of Akt and mTOR. However, this effect was weakened when PTEN was silenced. Furthermore, results in vivo showed that the expressions of PTEN, LC3 and Beclin-1 in chondrocytes of rats with KOA established by Hulth method treated with high dose DHJSD were significantly higher than those of the model group, while the expressions of p-Akt and p-mTOR were significantly lower than those of the model group. In addition, the formation of autophagosomes in chondrocytes of rats with KOA established by Hulth method treated with high dose DHJSD was significantly higher than that of the model group by transmission electron microscopy. So, it is reasonable to speculate that DHJSD can target PTEN to inhibit Akt/mTOR signaling pathway and promote the level of autophagy and the expression of LC3 and Beclin-1 proteins in chondrocytes.

As is well known, MMP-13 plays an important role in the progression of OA ^[33-35]^. Matrix metalloproteinases can be induced in multiple ways, and inflammatory cytokines such as IL-6 can directly induce production of matrix metalloproteinases, which is the pathological basis for the degradation of articular cartilage ^[36]^. This study showed that DHJSD directly inhibited the formation of IL-1β and MMP-13 *in vivo*, playing a protective role in articular cartilage.

Zinc is closely related to the formation of matrix metalloproteinases^[37, 38]^. In other words, when cells synthesize more matrix metalloproteinases in an inflammatory state, more zinc is needed to enter the cells. The present study showed that the intracellular zinc level increased after IL-1β treatment of chondrocytes, and the intracellular zinc level decreased after pretreatment with Ds in chondrocytes stimulated with IL-1β. Therefore, the downregulation of intracellular zinc level by DHJSD may be one of the reasons for inhibiting MMPs formation. So, DHJSD can enhance the level of autophagy in chondrocytes and LC3 expression, meanwhile reduce the level of zinc in chondrocytes. However, this study is not clear about how zinc in cells is excreted outside through autophagy and whether it is related to the cell exocytosis mechanism, which needs further confirmation (**Fig. 12**). Although this mechanism needs further clarification, it is not difficult to infer that the maintenance of intracellular zinc homeostasis by DHJSD is closely related to its enhancement of cellular autophagy.

**Fig. 12.**
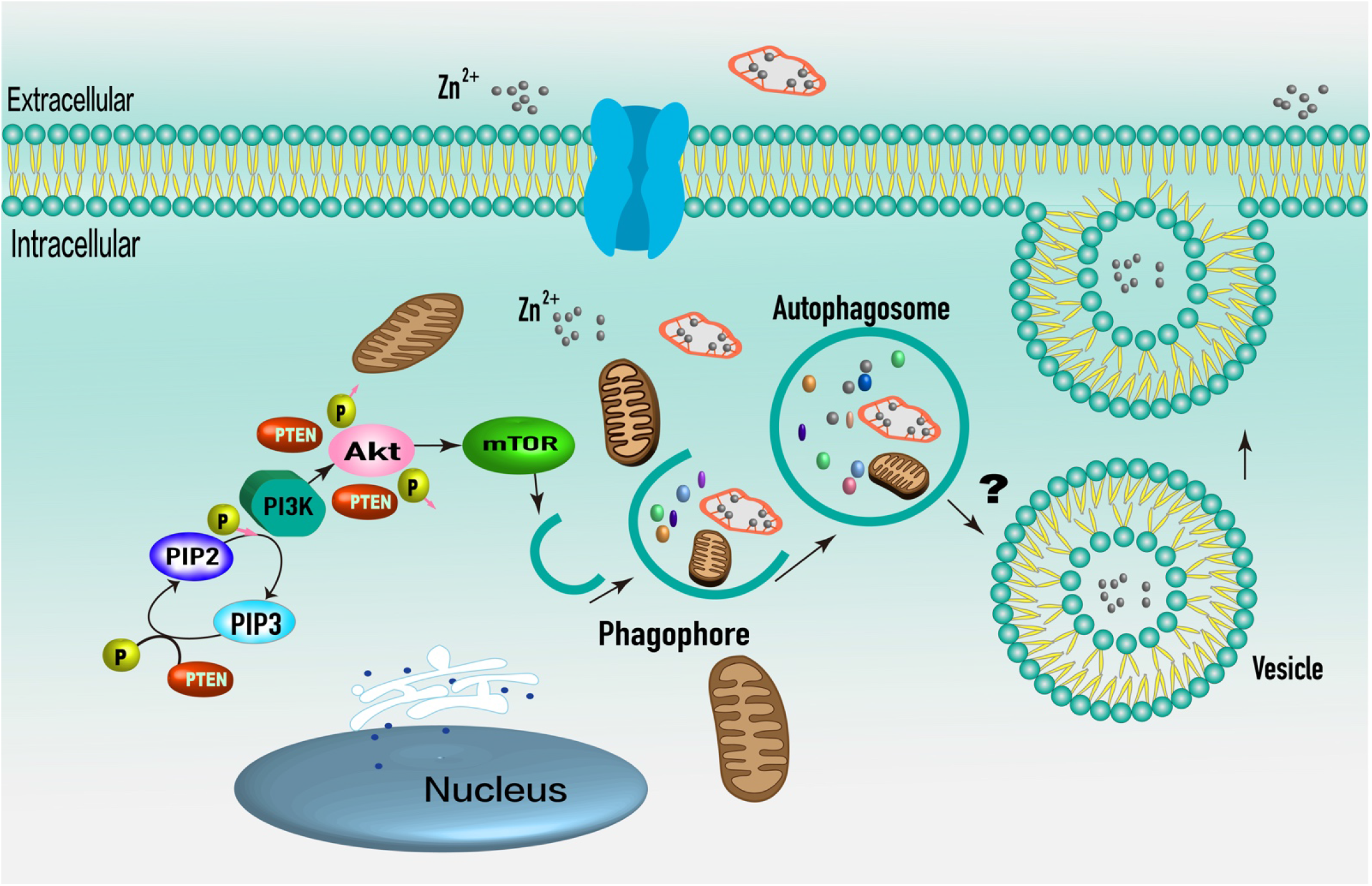
Schematic representation of the mechanism by which DHJSD regulates the zinc homeostasis in chondrocytes.

## 5. Conclusion

DHJSD can enhance the level of autophagy and regulate intracellular zinc homeostasis through PTEN/Akt/mTOR signaling pathway *in vivo*. In the rats with KOA triggered by Hulth method, DHJSD enhanced autophagy by inhibiting Akt/mTOR signaling pathway with PTEN, and down-regulated the proinflammatory cytokines IL-1β and MMP-13. In addition, histopathological observations verified the improvement of OA rats established by Hulth method treated with DHJSD. In conclusion, DHJSD has anti-osteoarthritis and cartilage protective effects in the treatment of OA *in vitro* and *in vivo*.

## Abbreviations

Akt: V-akt murine thymoma viral oncogene homolog
Bax: BCL2-Associated X
Bcl -2: B-cell lymphoma-2
DHJSD: Duhuo Jisheng Decoction
Ds: DHJSD-serum
HD: high dose
IL-1 β: interleukin-1β
ICP-MS/MS: inductively coupled plasma mass spectrometry
KOA: knee osteoarthri tis
LC-MS/MS: liquid chromatography tandem mass spectrometry
LC3: microtubule-associated prote ins 1A/1B light chain 3
LD: low dose
MD: medium dose
mTOR: mammalian target of rapamyci n
MMP-13: matrix metallopeptidase 13
OA: osteoarthritis
RT-PCR: real-time PCR
PTEN: phosph atase and tensin homolog
PI3K: phosphoinositide 3 kinase
p-Akt: phosphorylated Akt
p-mTOR: ph osphorylated mTOR.

## Author contributions

**Conceptualization**: Ye-Hui Wang

**Funding acquisition**: Ye-Hui Wang, Yi Zhou, Xiao-Hong Fan

**Methodology**: Ye-Hu Wang, Yi Zhou, Sheng Sun

**Project administration**: Ye-Hui Wang, Quan Xie

**Supervision**: Xiao-Hong Fan, Yang Fu

**Data curation**: Quan Xie, You-Peng Hu, Xiang Gao,

**Software**: Ye-Hui Wang, Yi-Zhou Xie

**Writing-original draft**: Ye-Hui Wang

## Acknowledgements

Thank professor Fan for writing-review and editing. Also, to the participants for their contributions in engagement as well as feedback.

## Data availability

Data will be made available on request.

## Declarations

### Ethics approval and consent to participate

The study was reviewed and approved by the Experimental Animal Ethics Committee of Hospital of Chengdu University of Traditional Chinese Medicine (No.2023DL-005). The experiment was carried out strictly accordance with the international rules and regulations of laboratory animal ethics. The study was carried out in compliance with the ARRIVE guidelines.

